# Evolution of influenza genome diversity during infection in immunocompetent patients

**DOI:** 10.1101/435263

**Authors:** Maxime Pichon, Bruno Simon, Martine Valette, Antonin Bal, Caroline Picard, Vanessa Escuret, Michèle Ottmann, Yves Gillet, Florence Ader, Bruno Lina, Laurence Josset

## Abstract

**Introduction:** Minor frequency viruses play many important roles during viral infection that cannot be explained by the consensus sequence alone. In influenza, immunosuppressed individuals appear to generate numerous viral variants, leading to subpopulations with important role in infection. The objective of the present study was to describe viral diversification over time in immunocompetent patients during influenza virus infection.

**Methods:** All clinical records of patients admitted to the Lyon university hospital (Lyon, France) during the influenza infection epidemics of the 2010-2015 period and sampled at least twice during their clinical management were retrospectively analyzed. To estimate performance of the sequencing procedures, well-characterized plasmids containing each of the 8 segments of influenza viruses were used as quality controls. Diversity, i.e. the number of validated single nucleotide variants, was analyzed to compare characteristics over time and according to clinical severity (mild, severe with neurological complications, severe with respiratory complications).

**Results:** After validation on quality controls (n=51), and verification of possible confusion bias, a 5%-threshold of detection was applied to clinical viral sequences (n=29). At this threshold, amino-acid coordinates (n=183/4,246, 4.31%) were identified as having at least one mutation during clinical course, independently of the clinical severity. Considering a threshold of 4 days of symptoms, as a limit for early and late sampling, diversity was significantly higher in late samples for the mild group, compared to both early mild and severe groups (p<0.05). At a single-segment scale, for PB2-coding segment, diversity was significantly higher in early samples of the neurological group than in both early and late samples in the respiratory group and for late samples in the mild group (p<0.05). For the NS1-coding segment, significant differences were observed between initial diversity of mild and severe patients, as for early and late samples in mild patients (p<0.01). Discussion. This study is the first describing diversity through time, associating biological and clinical information during viral diversification, during the infection of an immunocompetent human host. This latter opens a large field of investigation in infectious disease management using next-generation sequencing and suggest development of new therapies, focusing on non-antigenic viral properties, in non-vaccine fields of research

## Introduction

Influenza is a highly contagious respiratory viral infection for humans, mainly caused by influenza A (IAV) and B viruses (IBV), two of the five genera of the *Orthomyxoviridae* family. These viruses are enveloped viruses with a segmented, single-stranded and negative sense RNA genome (1). As for the other RNA viruses, genetic variability and plasticity of genomes are inherent properties of influenza viruses. Because of the lack of exonuclease activity and proof-reading function of viral RNA polymerase, replication of influenza virus genome has been evaluated to have a high mutation rate (more than 2 × 10^−5^ substitutions per nucleotide site per cell infection cycle) (2,3). Mutations are either rapidly lost in a non-advantageous environments (where they reduce the viral ability to replicate), or expanded (when they provide evolutionary advantage; for example antiviral resistance under selective pressure) (4). Because of the presence of minor frequency viruses, many important characteristics of a viral infection may not be entirely explained by the consensus sequence. Genetically diverse subpopulations co-circulating in the infected host, termed “viral quasispecies”, are generated upon replication of the infecting viruses (5). These viral quasispecies have been associated with rapid replication kinetics, and have many important implications for virus evolution and pathogenesis (6,7). In animal models, authors have demonstrated that a mutant poliovirus diverse population had more replicative and fitness advantages over its individual components, whereas high fidelity associated with lower diversity, generated strains unable to reach the brain of a susceptible infected mice (8). Some authors have proposed taking advantage of this mechanism for enhancement of vaccine security with an higher efficacy (8–10). Indeed, without quasispecies generation, cooperation between quasispecies is limited and vaccine strains could produce an improved immune response, with absence (or limited) adverse event. There is, however, little information for influenza disease. Some data have been obtained since the development of ultra-deep sequencing (UDS) through studies of infected individuals during the last pandemic or focusing on the hemagglutinin (HA) segment, or a broader study in immunosuppressed patients (11). This is of importance, as, similarly to other respiratory viruses (e.g. respiratory syncytial virus), immunosuppressed individuals appear to generate and spread numerous viral variants, and the selective pressure of antiviral therapy could lead to the emergence of antiviral-resistant viral variants (13,14). Although shedding can persist for months in the immunocompromised setting allowing full analysis of the viral diversification, the defective immune function enhances this trend (12). Thus, longitudinal data are required in immunocompetent patients, which was the objective of the present study aimed to describe viral diversification over time in human influenza virus infection.

## Methods

### Ethical statement

Respiratory samples (nasopharyngeal aspirate or swab) were collected for regular clinical management during hospital stay. No additional samples were taken for the purpose of this study. Patient confidentiality was strictly protected. This study was approved by the ethics committee of Hospices Civils de Lyon, France, on May 3, 2017.

### Patient selection

All clinical records of patients admitted to the Lyon university hospital (Hospices Civils de Lyon, France) during the influenza infection epidemics of the 2010-2015 period and who were sampled twice or more during their clinical management were retrospectively analyzed.

In order to standardize post-infection time, only patients who were symptomatic for 2 days or less when the first respiratory sample was collected were included in this study. With the aim to achieve a homogeneous panel of immunocompetent individuals and to limit potential confusion bias, those with at least one of the following were excluded: incomplete clinical file; having received antibiotics (antiviral, antibacterial, or antifungal) treatment; with a risk factor for severe influenza (list of criteria for inclusion in a vaccination plan, including chronic disease with or without treatment) (16). The clinical files of enrolled patients were retrospectively analyzed and patients were then classified according to clinical outcome. Patients were considered to have severe influenza when a respiratory and/or neurological complication(s) were reported (other were considered to have mild influenza). Those with severe influenza were further stratified according to complications. Neurological involvement was based on physician report of objective neurological abnormalities such as encephalopathy, encephalitis, focal symptoms or seizures and/or the necessity of admission in an intensive care unit (ICU) or in a specialized neurology unit (15). Respiratory failure criteria included patients with oxymetry <95% in an arterial sample and/or patients who required invasive or non-invasive ventilation and/or hospitalized in an ICU for respiratory distress.

Respiratory samples (NasoPharyngeal Aspiration – NPA, TracheoBronchial Aspiration – TBA, Broncho-Alveolar lavage – BAL, or Nasal Swabs – NS) from included patients were considered. All analyzed samples were tested positive for influenza A or B virus using Respiratory Multi Well System r-gene^®^ kit (bioMérieux, Marcy-l’étoile, France) during routine testing conducted at the virology department of the University Hospital of Lyon (allowing semi-quantification by determination of a Cycle threshold –Ct-). After routine screening, each specimen was stored at −20°C in Eagle’s Minimal Essential Medium with antibiotics. A limited number of freeze-thawing cycles were performed on ice, in order to conserve viral RNA.

Samples were considered as “early” or “late” sampling. Considering that a median time to symptoms alleviation is six days in a meta-analysis of trials focusing on benefit of antiviral treatment, early sampling was a day-1, day-2,day-3, and day-4 sampling and late sampling a day-5 or more sampling (17).

### Quality control preparation

To estimate performance of the wet- and dry- lab procedures, well-characterized pHW2000 plasmids containing each of the 8 segments of influenza viruses were used to constitute quality controls (18). Two different genetic backbones were used to simulate the two different IAV subtypes: A/Moscow/10/99 was used for H3N2, and A/Lyon/969/2009 was used for H1N1. To mimic viral diversification, mutated and non-mutated segments of one of the eight segments (*i.e.* the neuraminidase (NA) gene) were mixed at pre-defined concentrations. For H1N1, Three types of mutated NA plasmids, used for reverse genetic studies of the NA-resistance, were used (H275-mutated, M15I-mutated and V106I-N248D-N200S – triple mutated NA plasmid); for H3N2, a sole mutated plasmid was used (R292K-mutated NA). Each plasmid was quantified using QuBit fluorometer 2.0 (LifeTechnologies, Carlsbad, CA, USA) then pooled in determined concentration for each mutation to determine performances (0.1% for QC1; 0.5% for QC2; 1% for QC3; 5% for QC4; and 10% for QC5). A mix of non-mutated plasmid for both H1N1 and H3N2 subtypes was also used to verify the absence of false positive detection (QC0).

Mixes were then considered similarly to clinical samples for the rest of the study.

### Sample preparation

Nucleic acids were extracted from samples using an automatic extraction platform (Nuclisens EasyMag, bioMérieux). To optimize performance of subsequent following reverse transcription polymerase chain reaction (RT-PCR) and sequencing, without introducing to much bias, all samples were treated with DNAse. Briefly, 5μL of sample extract, 1μL of Turbo DNAse, 2μL of Turbo DNAse buffer (LifeTechnologies) and, to limit a potential residual RNAse activity, 0.5μL of RNasin^®^ Plus RNase Inhibitor (Promega Corporation, Madison, WI, USA) were incubated for 90 min at 37°C. Enzymatic reaction was then stopped by an immediate purification using a magnetic bead system (NucleoMag^®^ NGS Clean-up and Size Select, Macherey-Nagel, Düren, Germany) using a 0.5X ratio to limit potential small RNA presence.

Viral nucleic acids were amplified differentially depending on the viral type determined during routine testing. For IAV-positive samples, extracted nucleic acids were amplified using a multi-segment RT-PCR as described by Zhou *et al*. in 2009 (19). For IBV-positive samples, extracted nucleic acids were amplified using a RT-PCR as described by Zhou *et al.* in 2014 (20). Both of these protocol efficiencies were optimized using 0.5 μL of RNasin^®^ Plus RNase Inhibitor (Promega Corporation) to enhance PCR products quantities.

Correct amplification was verified by agarose gel electrophoresis (1%) and samples with no band visible in the gel were excluded from further analyses. Validated samples were purified using NucleoMag^®^ NGS Clean-up and Size Select (Macherey-Nagel) quantified first by Nanodrop (Thermo Fisher Scientific, Waltham, MA, USA) then by QuBit 2.0 fluorometer with HS DNA quantification kit (LifeTechnologies).

### Library preparation and sequencing

The library was constructed using the Nextera™ XT DNA library kit (Illumina, San Diego, CA, USA) according to manufacturer’s recommendations. Briefly, samples were fragmented and tagged during the same process by a Nextera XT transposase (Illumina). Then, these adaptor-ligated DNA fragments were linked to the barcode and adaptors combination by a limited-cycle PCR program (12 cycles). The resulting libraries were purified and size-selected using 0.5X NucleoMag^®^ beads (NucleoMag^®^ NGS Clean-up and Size Select, Macherey-Nagel). Using Nextera XT kit (Illumina), libraries were then pooled at an equimolar ratio, per 96, and then diluted in hybridation buffer. These multiplex samples were then denatured using NaOH and TrisHCl before loading on the NextSeq500 mid-output cartridge. After the 2×150 bp NextSeq paired-end sequencing run, data were base called and reads were collected and assigned to a sample when showing the same barcode combination, which generated Illumina FastQ files.

### Sequencing data

The reads obtained on the sequencing platform were submitted to NCBI’s Sequence Read Archive and can be found under project numbers SRP140895 (samples) and SRP137044 (quality control).

### Bioinformatic analysis

Briefly, the downstream analysis of the resulting Illumina FastQ files included: quality and read cleaning pre-processing, mapping to a generated reference and diversity analysis through Single Nucleotide Variants (SNV) calling (Figure 1). Adapters, low base-quality ends and read-pairs shorter than 50 bases were removed using cutadapt (v0.4.4) (21). Mapping to references was obtained using BWA-MEM alignment tool (v0.7.15) (22). Strain subtypes were verified on their hemagglutinin homology with subtype references. Consensus sequences were generated to obtain the closest possible mapping reference allowing a new mapping processing.

**Figure 1.**
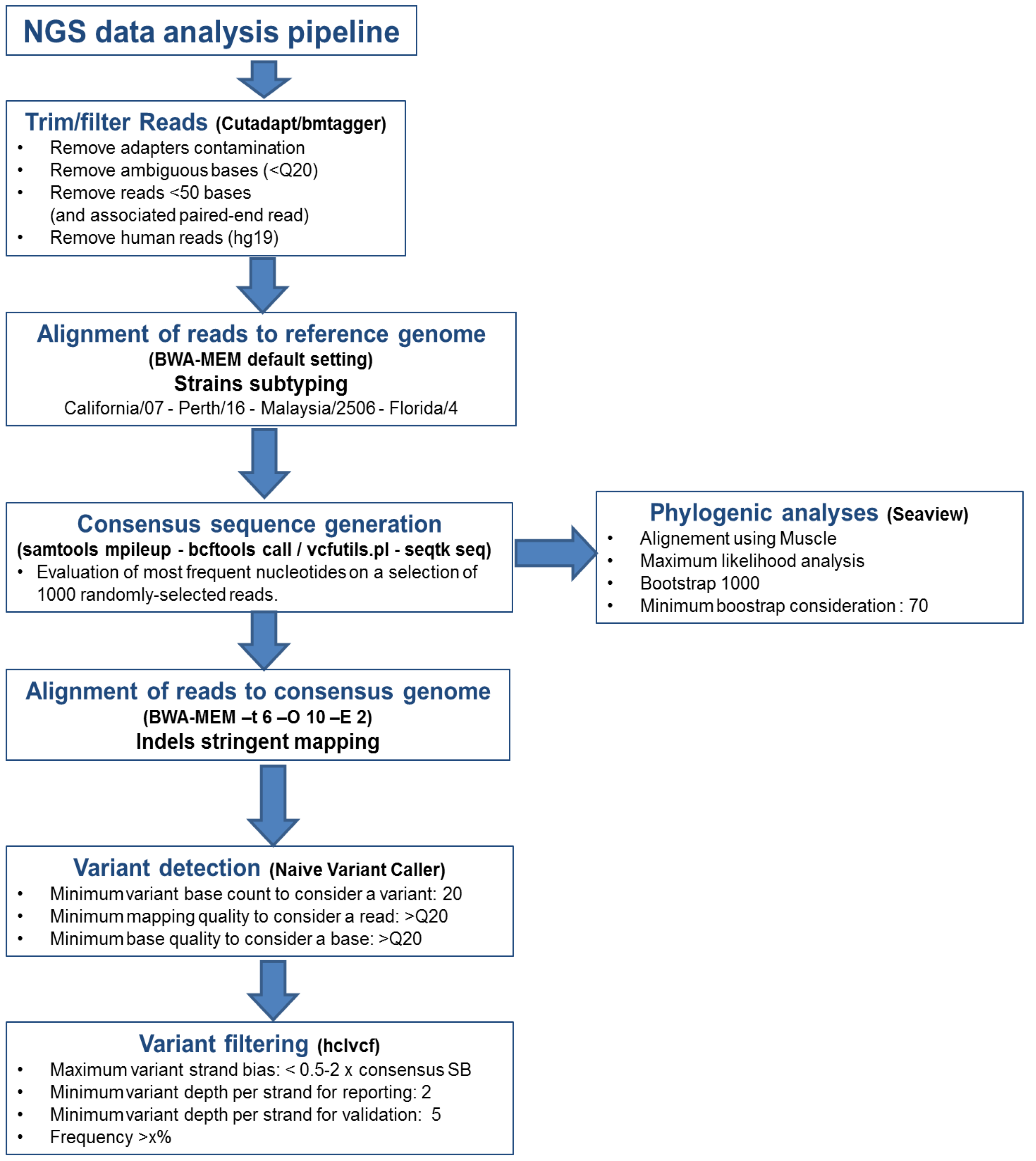
Sequencing data analysis pipeline. SB: strand-bias; Q20: Phred score of 20, hg19: human genome database 19^th^ version.

SNV calls were generated using naive variant caller (Biomina Galaxy platform) only filtering low quality bases (score<Q20) (23). End-user workable files, presenting validated SNVs and segment diversity, were generated using a homemade vcf analysis tool written in python (available upon request). Validation of these SNVs relied on stringent criteria determined on quality controls: variant’s strand bias < 1; minimal frequency estimated > 1%; frequency higher than a threshold based on the position coverage depth (24,25). Diversities, represented by the number of validated single nucleotide variant (mSNV), reported herein correspond to genomic polymorphism count weighted according to the proportion of the variants with respect to the main nucleotide.

FASTA consensus sequences were aligned using MUSCLE v3.8.31 (26). Trees, obtained using Maximum likelihood analyses on hemagglutinin (HA) sequences using IQ-Tree, were visualized and annotated using FigTree (v1.4.2) (27). Bootstrap values were estimated after 1,000 bootstrap replicates (28).

Statistical analyses and representations were performed using Graphpad Prism software (v7.0) (GraphPad Software Inc, La Jolla, CA, USA). Multiples testing were considered after Bonferroni's multiple comparison correction if needed. Results were considered to be significant when p-value (p) <0.05 (very significant when p<0.01)

## Results

### Selected patients and specimens

On the 692 clinical files screened for this study, 14 patients hospitalized from 2010 to 2015 with two or more respiratory samples (median number of samples per patient: 2; sampling delay between two samples from one to 14 days) tested positive for IAV or IBV were included for a total of 95 patient-days (i.e. a patient followed for a day). Clinical and biological data are summarized in Table 1. Ten patients were considered to have severe influenza (including seven in the respiratory group and three in the neurological group). Sex ratio was 1:1. Median age was 5 years (95%CI [1.7; 74.8]). Among these, ten had respiratory symptoms of variable severity (*i.e.* tachypnea, cough, rhinorrhea, bronchitis, wheezing) (10/14; 71.4%) and seven non-respiratory symptoms (*i.e.* diarrhea, vomiting, cutaneous eruption, adenopathy) (7/14; 50%). Over the clinical course of their disease, five patients (5/14; 35.7%; two patients in each of the severe groups and one in the mild group) received antivirals (oseltamivir treatment); nine patients received antibiotics, after sampling of the sequenced sample (3/3,100%, of those in the neurological group; 6/7, 85.7%, in the respiratory group). Three patients died (2/3, 66.6%, in the neurological group; 1/7; 14.3% in the respiratory group). All patients were sampled at least twice and one was sampled three times. Most of the 29 samples were taken from the upper respiratory tract (22 samples; 12 NS and 10 NPA); others were taken from the lower respiratory tract (7 samples; 6 BAL and 1 TBA).

**Table 1.** Demographic and clinical characteristics at baseline (A) and clinical evolution and therapeutic management (B) of all patients included in the study.

| Baseline demographic and clinical characteristics | Mild influenza<br>(n=4; 28.6%) | Severe influenza |  |
| --- | --- | --- | --- |
|  |  | Neurological<br>(n=3; 21.4%) | Respiratory<br>(n=7; 50.0%) |
| Baseline demographic characteristics |  |  |  |
| Sex, male (% of the group) | 4 (100) | 1 (33.3) | 3 (42.9) |
| Median age, years | 1.0 | 78.4 | 74.0 |
| Characteristics at the time of admission (% of the group) |  |  |  |
| First Sample Nature : NP aspirate | 3 (75) | 2 (66.6) | 1 (14.3) |
| First Sample Nature : NP swab | 1 (25) | 1 (33.3) | 6 (85.8) |
| Influenza B virus | 0 | 0 | 2 (28.6) |
| Influenza A virus (H1N1) | 1 (25) | 1 (33.3%) | 1 (14.3) |
| Influenza A virus (H3N2) | 3 (75) | 2 (66.7%) | 4 (57.2) |
| Comorbidities | 1* (25) | 0 | 0 |
| Clinical characteristics during clinical course (% of the group) |  |  |  |
| Ventilation | 0 | 2 (66.6) | 6 (85.7) |
| Signs of respiratory distress ** | 0 | 1 (33.3) | 6 (85.7) |
| Bacterial pneumonia complicating influenza | 0 | 2 (66.6) | 2 (28.6) |
| Coma | 0 | 1 (33.3) | 0 |
| Encephalitis |  | 2 (66.6) |  |
| Death | 0 | 2 (66.6) | 1 (14.3) |
| Treatment (% of the group) |  |  |  |
| No Antiviral use during clinical management (number of patients, % of the clinical group) | 0 | 1 (33.3%) | 5 (71.4%) |
| Antiviral use*** | 1 (25%) | 2 (66.6%) | 2 (28.6%) |
| No Antibiotic | 4 (100%) | 0 | 1 (14.3%) |
| Antibiotics | 0 | 3 (100%) | 6 (85.7%) |

\*Comorbidities: Non-malignant chronic anemia of unknown etiology (never associated with changes in viral diversity)
\*\* Signs of respiratory distress included: modification of breathing rate, increased heart rate, color changes, grunting, nose flaring, retractions, sweating wheezing, stridor, accessory muscle use, or changes in alertness.
\*\*\*Antiviral use: use of oseltamivir during the clinical course of the disease.

### Determination of SNV frequency threshold

A total of 51 artificial mixes of plasmids were sequenced to validate the process (27 H1N1-like plasmids and 24 H3N2-like plasmids). For each of sequencing run containing at least one sample sequenced for diversity analysis, at least one of these mixes of plasmids were sequenced. This allowed extensive performance validation of intra-run performance and inter-run reproducibility. Variants were detected in artificial mixes that included 10%, 5%, and 1% mutants (respectively named QC3, QC2, and QC1). The sequencing of a fully-wild type plasmid with a sufficient coverage (average depth 48,000X for H1N1-like plasmids and 44,000X for H3N2-like plasmids) found that there was no cross-contamination or bioinformatics bias (Table 2). Sufficient depth to allow statistical analyses was observed in clinical samples (Supplementary figure S1).

**Table 2.**
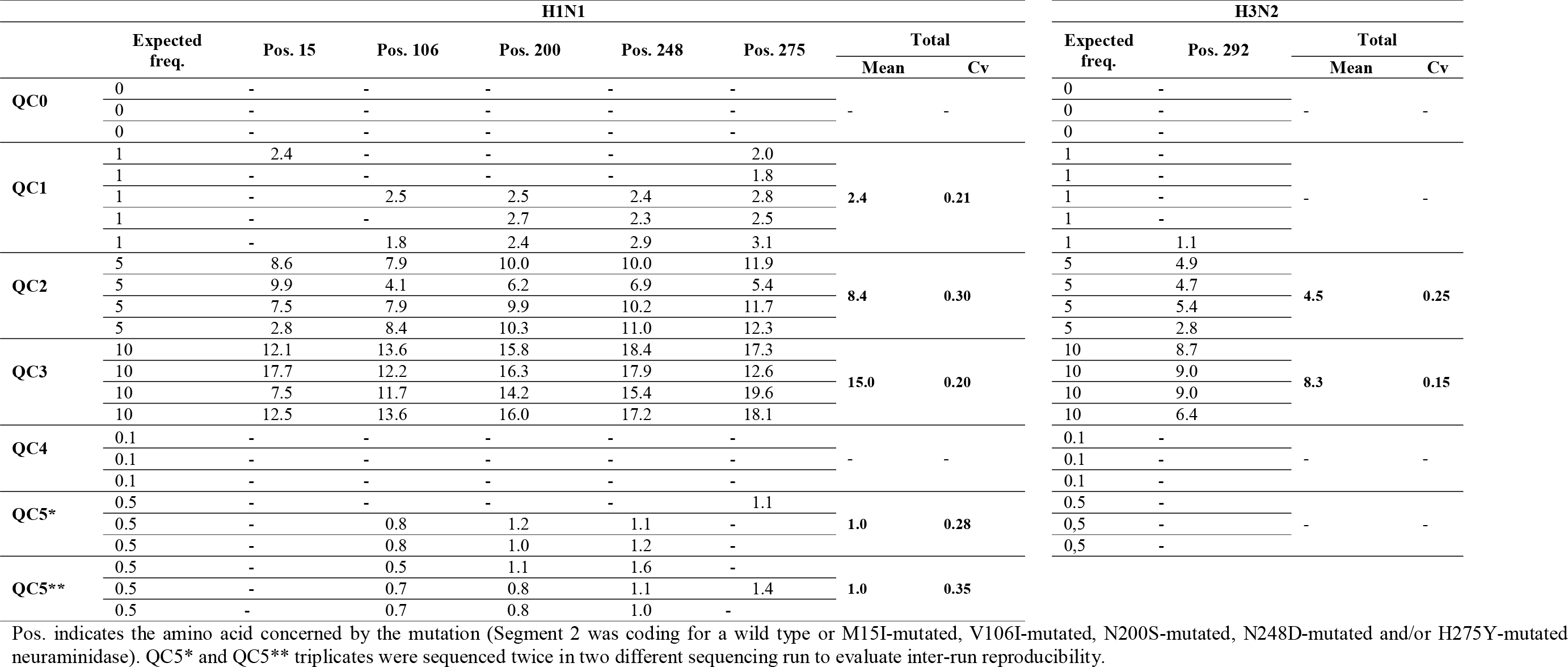
Quality Control (QC) performances.

All expected variants were detected for mixes containing 5% and 10% mutants (as summarized in Table 2). Sensitivity of detection decreased with the expected frequency of the mutation; resulting from inconstancies in detection for H1N1 (11 absence of detection on 25 nucleotide coordinates, 44%) and absence of detection in 80% of the sequenced QC for H3N2 at 1%-threshold. Mutants were not detected at 0.1% frequencies (named QC4) for both H1N1 and H3N2 plasmids and 0.5% (named QC5) for H1N1. Observed frequencies ranged from 150% to 250% of the expected frequency for H1N1; 82.8% to 89.2% for H3N2, without significant difference between mutations or replicates (p>0.05).

Moreover, to ensure good reproducibility of the sequencing process, and to evaluate possible inter-run bias, the plasmid mix with the lowest detected frequency (0.5% for H1N1-like plasmid) was sequenced in triplicate in two different sequencing runs. As there was no significant difference between the two runs (observed average frequencies 0.98 and 0.95, p>0.05), the grouped analysis of different runs was possible, considering sufficient inter-run reproducibility.

In light of these results, the detection threshold of 5% was used for the rest of the study.

### Influenza genome hotspots and coldspots

In IAV samples and using a 5% filter, 183 amino-acid coordinates (183/4,246, 4.31%) were identified as having at least one mutation during clinical course, independently of the clinical outcome (Table 3.). Prevalence of these residues represented less than 5% of all residue coordinates (from 3.16% for PA-coding segment to 6.61% for NA-coding segment, independently from oseltamivir treatment). There was no significant difference in viral number between segments (p>0.05).

As schematized in the Supplementary figure S2, hotspots (defined as regions with at least twice the mean mutation rate of the considered segment) were significantly more frequent in two of the eight major proteins (61/469 residues, 13.0% of the NA-coding segment; 21/229 residues, 9.17%, of the NS1-coding segment; *versus* 170/4246 residues, 4.00% for the whole influenza genome; p<0.05). Respective proportions or number of variants per segment were summarized in Figure 2. Finally, hotspots were less frequent in PB1- and M1-coding segments (24/757 residues, 3.17% of the total; 10/252 residues, 3.97% of the total segment *versus* 170/4246 residues, 4.00% for the whole influenza genome; p<0.05)

**Figure 2.**
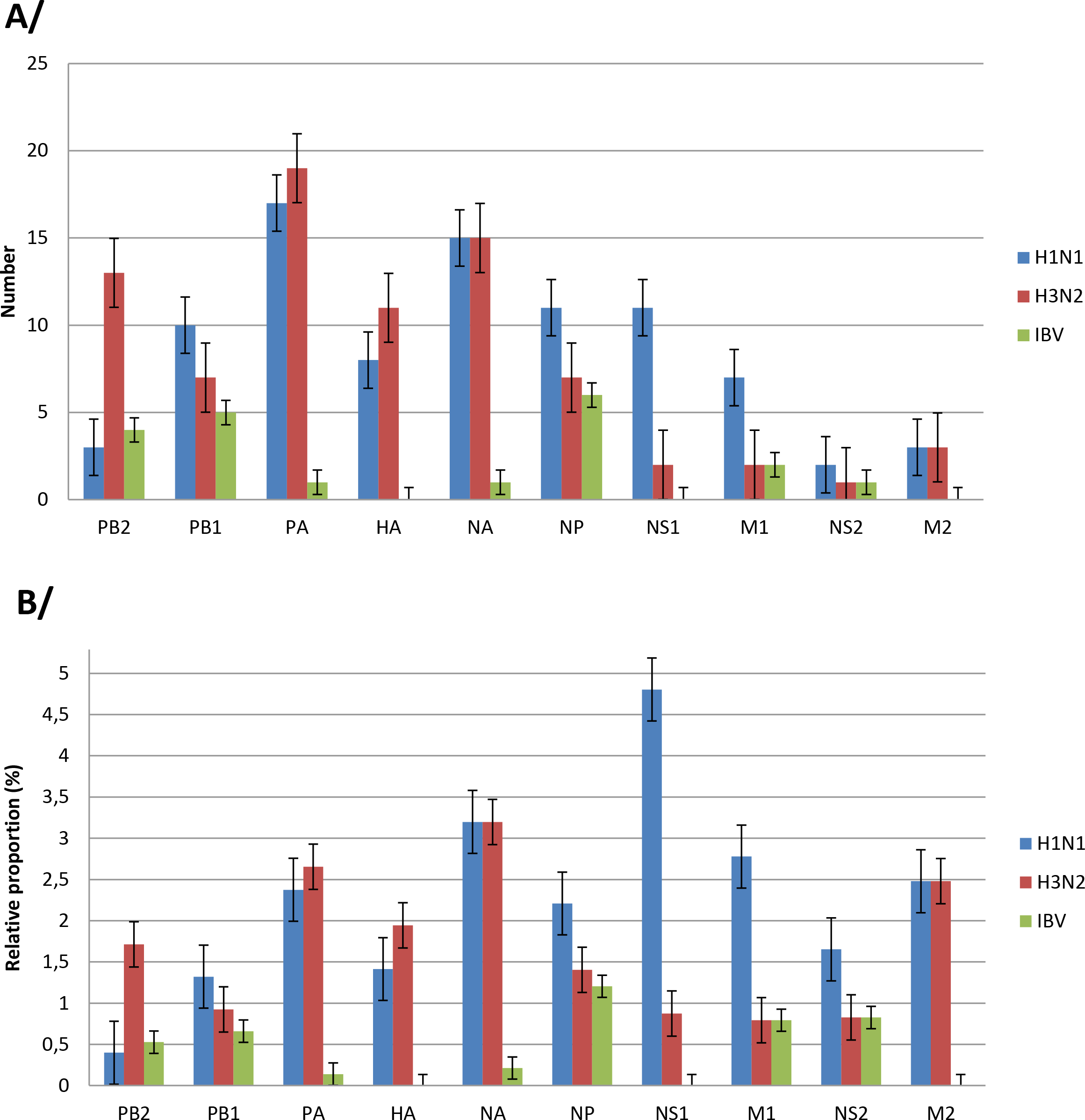
Average number (A) and relative proportions (B) of mSNV per segment per virus type/subtype. For each segment, three histograms were successively represented (H1N1 in blue, H3N2 in red and IBV in green). Error bar correspond to the standard error of numbers of mSNV (A) or mSNV frequencies (B). Only mSNV with frequency > 5% were considered (see material and methods).

### Intra-host diversity develops differentially according to disease outcomes

Even if samples could be clustered in the clade corresponding to their epidemiological season, no clustering could be observed in terms of clinical outcome or day of sampling since symptoms onset (p>0.05). Phylogenic analyses of the HA sequence of the most covered samples per patient is represented in Supplementary figure S3.

To overcome possible bias associated with the clinical non-standardized nature of sequenced samples, a statistical analysis between diversity and Ct was performed and did not find any significant association, allowing comparison of all samples (p>0.05).

Viral genomic diversity was evaluated in each patient, considering a threshold of 4 days of symptoms as a limit for early and late diversity. Diversity was found to significantly different between early and late samples (mean difference: 4.533, p<0.05; Figure 3a). Similarly, there were differences in frequencies when stratified according to time and clinical groups or subgroups (p<0.01). A significantly lower diversification was found in late samples of the mild group as compared to the severe neurological group (mean frequency difference 0.33, p<0.01) and as compared to the severe respiratory group (mean difference: 0.20, p<0.05; Figure 3b). A significantly lower diversity in late as compared to early samples of the mild group was also found (mean difference: 23.82, p<0.05). No significant correlation was found between diagnosis Ct and diversity observed after sequencing (p>0.05).

**Figure 3.**
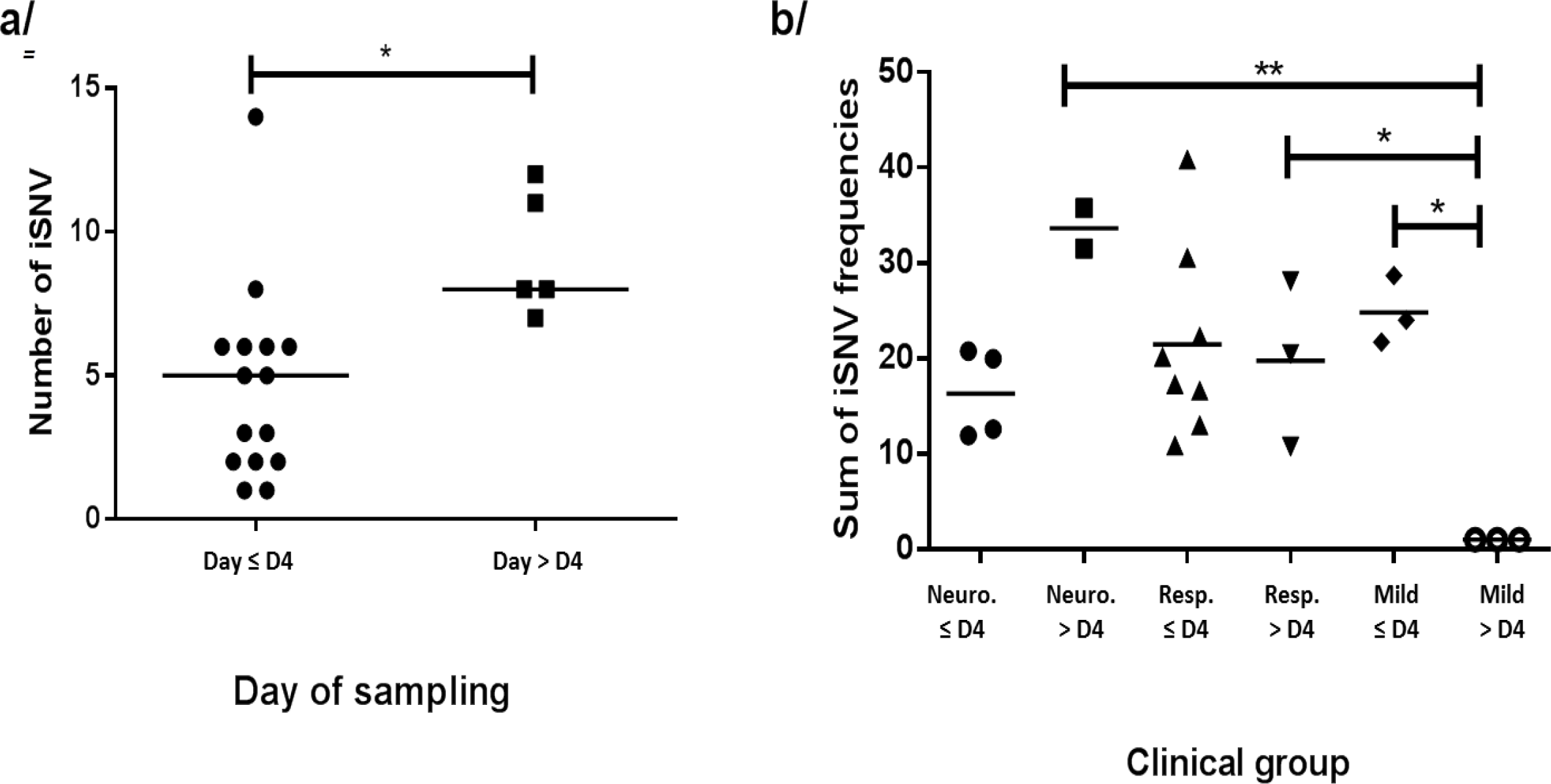
Comparison of intra-host diversity over time (whole genome). Diversity values (number or frequencies of Single Nucleotide Variants – mSNV – observed in a sample, summarizing the diversity of all segments) are represented according to the time of sampling (before or after four days of symptoms, as no specimen was sampled at Day 4). * p < 0.05. ** p < 0.01.

Analyses were then performed at a single-segment scale (significant results were summarized in Figure 4). Diversity was significantly lower in early than in late samples (mean difference: 2.55; p<0.05) for segment 3 (PB2-coding segment; Figure 4a). For this segment, there was significantly greater diversity in the neurological group in early samples than in both early and late samples in the respiratory groups (mean differences: 6.44 and 7.33, respectively; p<0.05) and for late samples in the mild group (mean difference 8.00; p<0.05, Figure 4b). On the NS1-coding segment, diversity was not significantly different in early and in late samples for segment 8 (Figure 4c). On this segment, significant differences were observed between diversity in specimens sampled early in the clinical course of mild-group patients and of severe-group patients (mean difference 5.00 and 6.08; p<0.05 and p<0.01 for comparison with respiratory and neurological groups respectively). Moreover, a very significant difference was observed between early and late diversity for mild-group patients (mean difference 6.33; p<0.01; Figure 4d). Analyses performed using a 1%-threshold for SNV calling found similar results.

**Figure 4.**
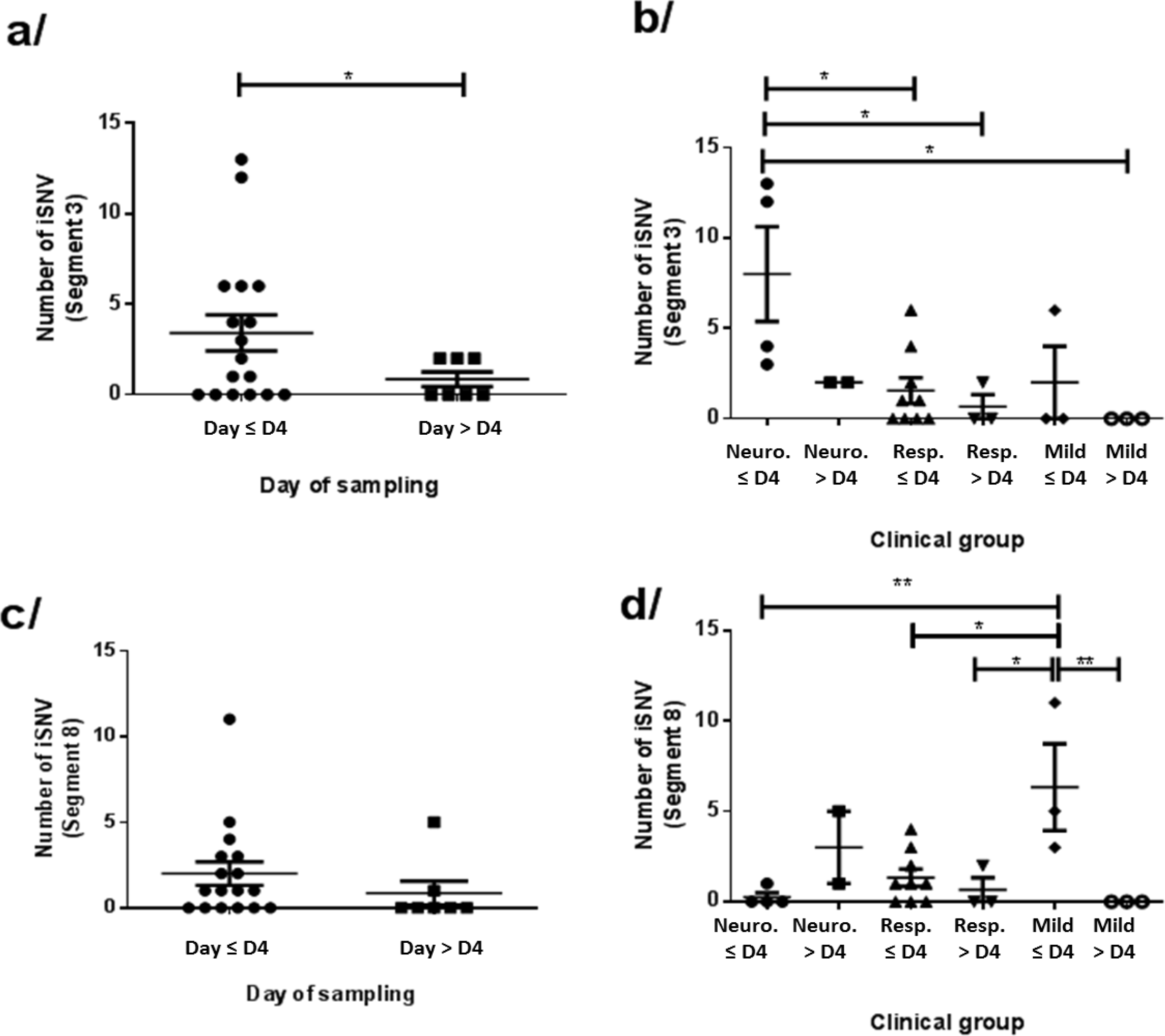
Comparison of intra-host diversity over time (per segment). Diversity values (number or frequencies of Single Nucleotide Variants – SNV – observed in a specific segment of influenza virus) for (a and b) Segment 3 (or PB2) and (c and d) Segment 8 (or NS) are represented in function of the time of sampling (before or after four days of symptoms). * p <0.05. ** p < 0.01. Note that to ease understanding, only segment with statistical differences between early and late samples were represented on this figure.

## Discussion

This study is the first, to the best of our knowledge, to describe influenza virus diversification over time using clinical specimens sampled in immunocompetent patients and ultra-deep sequencing.

Before the emergence of UDS, “classical” Sanger sequencing did not allow deep characterization of these viral populations, due to insufficient depth of sequencing. At contrary, UDS seems to fulfill this requirement (29–31). However, owing to possible experimental errors, introduced both during wet-lab (*i.e.* RT-PCR but also sequencing itself) and dry-lab (*i.e.* bioinformatic analyses) steps, appropriately prepared quality controls are required to valid these processes (32). As UDS enables sequencing of several gigabases of DNA in a single run, and because influenza genome is a limited 13 kb-long genome, a single sequencing could allow its characterization with a high coverage (*i.e.* the proportion of the genome that is sufficiently sequenced) and/or depth (*i.e.* the number of times a considered nucleotide coordinate is sequenced on a reference genome) (33). Nevertheless, the segmented nature of its genome makes it technically difficult to obtain sufficient (and homogenous) coverage of all eight genomic segments in a single reaction to allow diversity analyses. The present study describes the use of adapted quality controls to validate the comparison between runs and between samples in viral variant analysis. Using well-characterized plasmids, all expected mutations were observed using a frequency threshold of 5% for SNV variant calling. Using only two different plasmids, a similar approach defined a frequency threshold of 2%, but as described by the authors the pipeline used does not take strand bias into account, justifying the supplementary precaution taken herein (even if no supplementary difference was observed using a threshold of 1%) (25,34). Furthermore, plasmids are a more accurate method to evaluate performance of NGS than whole viruses isolated in plaque assays as these contain confounding mutations introduced during replication (35). It is important to note that, herein, observed differences remained the same independently from the frequency filter that was applied. Because the frequency threshold chosen has to be inversely related to the depth of sequencing, this observation suggests that data from less deep sequencing remain of interest to compare samples sequenced using the same method, which is of interest for future studies.

In the present study, respiratory specimens were limited to those sampled from immunocompetent patients before the introduction of antimicrobial therapy which allowed the description of a baseline state of viral diversity without identifiable confounding factor. This baseline diversity has already been described as resulting from a bottleneck in studies investigating other types of virus (36). First, we observed that the complete genome scale diversity increases during clinical course. Secondly, there were differences in terms of genomic diversification according to clinical outcome; while a significant decrease was found between early and late diversity in patients with mild influenza, no significant difference was found for those with severe disease either in the neurological or respiratory group. This difference suggests that diversification could be considered as a virulence factor for influenza, as has been suggested for Coxsackie virus, and as described in a poliovirus mouse model, and for Foot-and-Mouth Disease virus (8,37,38). Moreover, this result also supports the potential value of an engineered virus with high fidelity polymerase to improve vaccine effectiveness (10). Indeed, responsible for less severe manifestations, less diversified viruses obtained using these polymerases would enhance vaccine effectiveness without a dramatic increase in morbidity rates. Conversely, the dramatic decrease of diversity observed for mild influenza is consistent with the notion of an error catastrophe threshold, as previously suggested in therapeutic interventions (39). The observed results in the present study could suggest that a decrease of diversity, associated with a viral clearance and improvement of symptoms, reflects that the error threshold was exceeded that led to extinction of the viral population. This observation could support the value of ribavirin, an antiviral drug that destabilizes the polymerase, in influenza infection, as has been reported for Haanta virus (40). Furthermore, the attenuated virulence of less diverse influenza virus populations could support the use bioengineered virus bearing high-fidelity polymerase to produce less diverse quasispecies and therefore enhance vaccine effectiveness without risk of secondary effects.

At the single segment scale, genomic diversity evolved differentially according to the considered segment. The significant decrease observed at the level of the whole genome between early and late samples was driven by evolutions of PB2-coding and NS1-coding segments. These differences in diversity were associated with the clinical groups, with a higher initial diversity in the severe neurological influenza group compared to both respiratory and mild clinical groups in the PB2-coding segment, at the converse in the NS1-coding segment.

It has previously been reported that NS1-coding segment mutations could be associated with clinical outcome, for instance in equine model of infection or in highly pathogenic influenza H5N1 infections (41,42). Non-antigenic segments (all but HA- and NA-coding segments) could also be considered as a fitness-enhancing capacity for viruses, as demonstrated (43). For example, substitutions in the nucleoprotein were considered as to play a major role in adaptation and virulence, enhancing the capacity to escape to T cell mediated immunity (44). The present study analyzed the diversity per nucleotide coordinate as a whole and not only mSNV, and the data presented suggest that segments carrying high diversity could be associated with clinical outcome, bringing potential properties to newly-created viruses.

The present study suffers from some limitations, mainly due to its retrospective nature. The most important is the limited size that reduces the precision of observed SNV at a single nucleotide coordinate level, but this did not impact greatly the results obtained. Moreover, the sampling time points were those of routine care, and for research purposes it would be of interest to increase the sampling frequency in order to obtain a finer analysis of daily diversity variations. This analysis could validate the hypothesis of error-catastrophe threshold and determine the level of this, in order to transfer these notions to clinical management. However, without description of the clinical outcome, it has already been reported that variation over less than one day could be considered as negligible, representing few viral replication cycle.

Nevertheless, this work remains a preliminary study for influenza virus genomic diversification, associating biological and clinical information during viral diversification, during the infection of a human host. This latter opens a large field of investigation in infectious disease management using next-generation sequencing and suggest development of new therapies, focusing on non-antigenic viral properties, in non-vaccine fields of research.

## Acknowledgments

We thank Gwendolyne Burfin (CNR Respiratory virus), for her technical assistance as well as Philip Robinson (DRCI, Hospices Civils de Lyon) for his valuable help in manuscript preparation.

## Author contributions

**MP**: performed the sample preparations and sequencing, and performed bioinformatic analysis, writing - original draft; main writer of the manuscript; **BS**: performed bioinformatic analysis, writing - original draft; **MV**: performed the sample preparations, provided the QC system; **AB**: performed the sample preparations; **VE**: guarantor for clinical data; **MO**: provided the QC system; **EJ**: guarantor for clinical data; **YG**: guarantor for clinical data; **BL**: guarantor for clinical data; **LJ**: conceptualization, formal analysis, writing - original draft, guarantor for the NGS data. All authors reviewed and approved the final version of the manuscript.

## Supplementary material

Supplementary figure S1. Coverage plot for samples

Supplementary figure S2. Consensus minor variants contextualized within influenza genome segments. Each of the ten major proteins encoded by each of the influenza genome segments (major proteins: PA, PB1, PB2, HA, NA, NP, NS1, M2, an alternative proteins: NS2, M2 are represented). On this representation, substitutions vanishing or emerging between first and second sample for each patient are indicated in red and green respectively (in yellow are represented coordinates with both indication, depending of the considered sample). Orange dots correspond to residues implicated in a hotspot and light blue dots to those implicated in coldspot regions.

Supplementary figure S3. Phylogenic analyses of sequenced samples.

## References

1. Wright P, Webster R. Orthomyxoviruses. In: Knipe D, Howley P, editors. Fields Virology. Philadelphia: Wilkins; 2001.

2. Nobusawa E, Sato K. Comparison of the mutation rates of human influenza A and B viruses. J Virol. 2006;80(7):3675–8.

3. Sanjuan R, Nebot MR, Chirico N, Mansky LM, Belshaw R. Viral Mutation Rates. J Virol. 2010 Oct 1;84(19):9733–48.

4. Lauring AS, Andino R. Quasispecies theory and the behavior of RNA viruses. PLoS Pathog. 2010;6.

5. Domingo E, Sheldon J, Perales C. Viral quasispecies evolution. Microbiol Mol Biol Rev. 2012;76(2):159–216.

6. Ramakrishnan MA, Tu ZJ, Singh S, Chockalingam AK, Gramer MR, Wang P, et al. The feasibility of using high resolution genome sequencing of influenza A viruses to detect mixed infections and quasispecies. PloS One. 2009;4(9):e7105.

7. Stech J, Xiong X, Scholtissek C, Webster RG. Independence of evolutionary and mutational rates after transmission of avian influenza viruses to swine. J Virol. 1999 Mar;73(3):1878–84.

8. Vignuzzi M, Stone JK, Arnold JJ, Cameron CE, Andino R. Quasispecies diversity determines pathogenesis through cooperative interactions in a viral population. Nature. 2006;439.

9. Pfeiffer JK, Kirkegaard K. Increased fidelity reduces poliovirus fitness and virulence under selective pressure in mice. PLoS Pathog. 2005 Oct;1(2):e11.

10. Vignuzzi M, Wendt E, Andino R. Engineering attenuated virus vaccines by controlling replication fidelity. Nat Med. 2008 Feb;14(2):154–61.

11. Pichon M, Picard C, Simon B, Gaymard A, Renard C, Massenavette B, et al. Clinical management and viral genomic diversity analysis of a child’s influenza A(H1N1)pdm09 infection in the context of a severe combined immunodeficiency. Antiviral Research. (Submitted);

12. Baz M, Abed Y, McDonald J, Boivin G. Characterization of multidrug-resistant influenza A/H3N2 viruses shed during 1 year by an immunocompromised child. Clin Infect Dis. 2006;43(12):1555–61.

13. Geis S, Prifert C, Weissbrich B, Lehners N, Egerer G, Eisenbach C, et al. Molecular characterization of a respiratory syncytial virus outbreak in a hematology unit in Heidelberg, Germany. J Clin Microbiol. 2013;51(1):155–62.

14. van der Vries E, Stittelaar KJ, van Amerongen G, Veldhuis Kroeze E, de Waal L, Fraaij PLA, et al. Prolonged influenza virus shedding and emergence of antiviral resistance in immunocompromised patients and ferrets. PLoS Pathog. 2013;9(5):e1003343.

15. Khandaker G, Zurynski Y, Buttery J, Marshall H, Richmond PC, Dale RC, et al. Neurologic complications of influenza A(H1N1)pdm09: Surveillance in 6 pediatric hospitals. Neurology. 2012;79(14):1474–81.

16. Gill PJ, Ashdown HF, Wang K, Heneghan C, Roberts NW, Harnden A, et al. Identification of children at risk of influenza-related complications in primary and ambulatory care: a systematic review and meta-analysis. Lancet Respir Med. 2015;3(2):139–49.

17. Dobson J, Whitley RJ, Pocock S, Monto AS. Oseltamivir treatment for influenza in adults: a meta-analysis of randomised controlled trials. Lancet. 2015;385(9979):1729–37.

18. Hoffmann E, Neumann G, Kawaoka Y, Hobom G, Webster RG. A DNA transfection system for generation of influenza A virus from eight plasmids. PNAS USA. 2000;97(11):6108–13.

19. Zhou B, Donnelly ME, Scholes DT, St George K, Hatta M, Kawaoka Y. Single-reaction genomic amplification accelerates sequencing and vaccine production for classical and Swine origin human influenza A viruses. J Virol. 2009;83.

20. Zhou B, Lin X, Wang W, Halpin RA, Bera J, Stockwell TB. Universal influenza B virus genomic amplification facilitates sequencing, diagnostics, and reverse genetics. J Clin Microbiol. 2014;52.

21. Martin M. Cutadapt removes adapter sequences from high-throughput sequencing reads. EMBnet.journal. 2011;17(1):10.

22. Li H, Handsaker B, Wysoker A, Fennell T, Ruan J, Homer N, et al. The Sequence Alignment/Map format and SAMtools. Bioinforma Oxf Engl. 2009;25(16):2078–9.

23. Blankenberg D, Von Kuster G, Bouvier E, Baker D, Afgan E, Stoler N, et al. Dissemination of scientific software with Galaxy ToolShed. Genome Biol. 2014;15(2):403.

24. Guo Y, Cai Q, Samuels DC, Ye F, Long J, Li C-I, et al. The use of next generation sequencing technology to study the effect of radiation therapy on mitochondrial DNA mutation. Mutat Res Toxicol Environ Mutagen. 2012;744(2):154–60.

25. Guo Y, Li J, Li C-I, Long J, Samuels DC, Shyr Y. The effect of strand bias in Illumina short-read sequencing data. BMC Genomics. 2012;13(1):666.

26. Edgar RC. MUSCLE: multiple sequence alignment with high accuracy and high throughput. Nucleic Acids Res. 2004;32(5):1792–7.

27. Nguyen L-T, Schmidt HA, von Haeseler A, Minh BQ. IQ-TREE: a fast and effective stochastic algorithm for estimating maximum-likelihood phylogenies. Mol Biol Evol. 2015;32(1):268–74.

28. Minh BQ, Nguyen MAT, von Haeseler A. Ultrafast Approximation for Phylogenetic Bootstrap. Mol Biol Evol. 2013 May 1;30(5):1188–95.

29. Junemann S, Sedlazeck FJ, Prior K, Albersmeier A, John U, Kalinowski J. Updating benchtop sequencing performance comparison. Nat Biotechnol [Internet]. 2013;31—. Available from: http://dx.doi.org/10.1038/nbt.2522

30. Loman NJ, Misra RV, Dallman TJ, Constantinidou C, Gharbia SE, Wain J. Performance comparison of benchtop high-throughput sequencing platforms. Nat Biotechnol. 2012;30.

31. Quail MA, Smith M, Coupland P, Otto TD, Harris SR, Connor TR. A tale of three next generation sequencing platforms: comparison of Ion Torrent. Pacific Biosciences and Illumina MiSeq sequencers. BMC Genomics. 2012;13.

32. Robasky K, Lewis NE, Church GM. The role of replicates for error mitigation in next-generation sequencing. Nat Rev Genet. 2014;15.

33. Glenn TC. Field guide to next-generation DNA sequencers. Mol Ecol Resour [Internet]. 2011;11. Available from: http://dx.doi.org/10.1111/j.1755-0998.2011.03024.x

34. McCrone JT, Woods RJ, Martin ET, Malosh RE, Monto AS, Lauring AS. Stochastic processes constrain the within and between host evolution of influenza virus. eLife. 2018;7.

35. Dulbecco R, Vogt M. Plaque formation and isolation of pure lines with poliomyelitis viruses. J Exp Med. 1954 Feb;99(2):167–82.

36. McCrone JT, Lauring AS. Genetic bottlenecks in intraspecies virus transmission. Curr Opin Virol. 2018;28:20–5.

37. Moratorio G, Henningsson R, Barbezange C, Carrau L, Bordería AV, Blanc H, et al. Attenuation of RNA viruses by redirecting their evolution in sequence space. Nat Microbiol. 2017;2:17088.

38. Rai DK, Diaz-San Segundo F, Campagnola G, Keith A, Schafer EA, Kloc A, et al. Attenuation of Foot-and-Mouth Disease Virus by Engineered Viral Polymerase Fidelity. J Virol. 2017;91(15).

39. Mori K, Murano K, Ohniwa RL, Kawaguchi A, Nagata K. Oseltamivir expands quasispecies of influenza virus through cell-to-cell transmission. Sci Rep. 2015;5:9163.

40. Severson WE, Schmaljohn CS, Javadian A, Jonsson CB. Ribavirin causes error catastrophe during Hantaan virus replication. J Virol. 2003;77(1):481–8.

41. Chambers TM, Quinlivan M, Sturgill T, Cullinane A, Horohov DW, Zamarin D, et al. Influenza A viruses with truncated NS1 as modified live virus vaccines: pilot studies of safety and efficacy in horses. Equine Vet J. 2009 Jan;41(1):87–92.

42. Steel J, Lowen AC, Pena L, Angel M, Solórzano A, Albrecht R, et al. Live attenuated influenza viruses containing NS1 truncations as vaccine candidates against H5N1 highly pathogenic avian influenza. J Virol. 2009;83(4):1742–53.

43. Raghwani J, Thompson RN, Koelle K. Selection on non-antigenic gene segments of seasonal influenza A virus and its impact on adaptive evolution. Virus Evol. 2017;3(2):vex034.

44. Gong L., Bloom JD. Epistatically interacting substitutions are enriched during adaptive protein evolution. PLoS Genet. 2014;10(5):e1004328.

